# Domain Binding and Isotype Dictate the Activity of Anti-human OX40 Antibodies

**DOI:** 10.1101/2020.06.29.177691

**Authors:** Jordana Griffiths, Khiyam Hussain, Hannah L Smith, Theodore Sanders, Kerry Cox, Monika Semmrich, Linda Martensson, Jinny Kim, Tatyana Inzhelevskaya, Chris Penfold, Alison L Tutt, Ian M Mockridge, Claude HT Chan, Vikki English, Ruth R French, Ingrid Teige, Aymen Al-Shamkhani, Martin J Glennie, Bjorn Frendeus, Jane E Willoughby, Mark S Cragg

**Affiliations:** Antibody and Vaccine Group, Centre for Cancer Immunology, Cancer Sciences Unit, Faculty of Medicine, University of Southampton, Tremona Road, Southampton SO16 6YD, UK; Preclinical Research, BioInvent International AB, Sölvegatan 41, 22370 Lund, Sweden

**Keywords:** OX40, CD134, Fc gamma receptor, antibody immunotherapy, isotype, Treg T cell expansion, tumour immunotherapy

## Abstract

Previous data suggests that anti-OX40 mAb can elicit anti-tumour effects in mice through deletion of Tregs. However, OX40 also has powerful costimulatory effects on T cells which could evoke therapeutic responses. The contributions of these different effector mechanisms has not previously been systematically evaluated, particularly for mAb directed to human OX40. Therefore, we generated a novel human OX40 knock-in (KI) mouse to evaluate a panel of anti-hOX40 mAb and show that their activities relate directly to two key properties: 1) isotype – with mIgG1 mAb evoking receptor agonism and CD8+ T cell expansion and mIgG2a mAb evoking deletion of Treg and; 2) epitope - with membrane-proximal mAb delivering more powerful agonism. Intriguingly, both isotypes acted therapeutically in tumour models by engaging these different mechanisms. These findings highlight the significant impact of isotype and epitope on the modulation of anti-hOX40 mAb therapy, and indicate that CD8+ T cell expansion or Treg depletion might be preferable according to the composition of different tumours.

**Summary:** Human trials with anti-OX40 antibodies have been disappointing indicating that optimal reagents have not yet been developed. Here, using a new panel of antibodies, we show that isotype and epitope combine to determine agonistic and therapeutic activity.

## Introduction

The use of immunomodulating monoclonal antibodies (mAb) to generate anti-tumour immune responses offers an exciting approach to cancer immunotherapy. mAb against immune checkpoint inhibitors such as Ipilimumab and Nivolumab, which target the co-inhibitory receptors CTLA-4 and PD-1, respectively, pioneered this approach and have demonstrated success in treating a number of previously untreatable cancers [1, 2]. However, many patients do not respond to these reagents and additional therapeutic strategies are required. Agonistic mAb targeting co-stimulatory receptors have emerged as targets for clinical development, in particular members of the tumour necrosis family of receptors (TNFR) such as CD40 [3], 4-1BB [4], GITR [5], OX40 [6–8], and CD27 [7, 9]. However a recent paper by Freeman et al identified a intratumoural Treg signature which included TNFR family members with the hypothesis that they could be targeted instead by depleting antibodies in order to generate therapy [10]. TNFR family members are typically characterised by an extracellular domain (ECD) consisting of several cysteine rich domains (CRDs) which allow for binding of their respective trimeric ligands leading to receptor clustering and downstream signalling [11]. mAb targeting such receptors have been shown to depend on their interaction with the inhibitory FcγR (FcγRIIB) to generate sufficient cross-linking and resultant agonistic activity [12, 13]. More recently, however, the ability of a number of TNFR and CTLA-4 targeting mAb have been shown to cause deletion of Tregs via engagement of activatory FcγR [14–17]. One example is the anti-mouse OX40 mAb, OX86 (rat IgG1), which for many years was known to enhance effector T cell proliferation and survival leading to successful therapeutic outcomes in pre-clinical models [6, 18]. However, it was recently also shown to be capable of deleting Tregs in an activatory FcγR-dependent manner [14]. This effect was directly influenced by isotype, with mIgG2a showing greater depleting capacity than the native rIgG1 isotype. mOX40 expression levels were also shown to be important with preferential depletion of Tregs over effector T cells being due to differential OX40 expression on these T cell subsets [14].

Previous work studying the mechanisms of action of therapeutic antibodies has shown that the region of the molecule targeted is also an important consideration, with even subtle shifts in binding epitope resulting in large impacts on effector function and efficacy [19, 20]. Recent work on several TNFRs has further highlighted the importance of these aspects in influencing the type and strength of effector function [20–23]. For example, we demonstrated the importance of domain binding on the biological activity of anti-CD40 mAb, showing that membrane distal CRD1-binding mAb were strong agonists of CD40 with membrane proximal mAb less potent [22]. Furthermore, mAb binding CRD2-4 blocked CD40L and were potent antagonists. We also demonstrated a domain binding association with differential effector mechanisms for anti-4-1BB mAb [20], showing mAb which bound membrane proximal domains engaged in more effective CDC and ADCC killing mechanisms with ADCP less affected [20]. Recently, Zhang et al. reported that mAb binding to mouse (m)OX40 which blocked ligand binding and bound CRD2, or bound at the membrane proximal domain (CRD4), provide stronger agonistic and anti-tumour activity than mAb binding CRD1 and 3 [23]. These results differed from those seen for hCD40, highlighting that the functional effects of mAb domain binding are likely to require assessment for each of the TNFR family members and validation for each species.

Given these discrepancies, we explored the optimal domain binding and isotype for a novel panel of anti-human (h)OX40 mAb which collectively bound to all 4 CRDs of the ECD of hOX40. We evaluated their function *in vitro* and *in vivo* as both mIgG1 and mIgG2a isotypes. Using a novel hOX40 knock-in (KI) mouse we found that mIgG1 mAb were agonistic and engendered memory responses, whereas mIgG2a mAb had depleting activity with poorer memory recall to a challenge antigen. We observed that the strength of these effector functions correlated with domain binding; those mAb which bound to the most membrane proximal domain (CRD4) which did not block ligand binding, showed the strongest agonistic activity as mIgG1 as well as the most potent depleting activity as mIgG2a. This data highlights how mAb to different TNFR (and different species) exhibit different requirements in relation to optimal domain binding and effector function and indicate how more active anti-hOX40 mAb might be developed.

## Results

### hOX40KI mice express hOX40 and develop normally

In order to investigate the immunotherapeutic potential of a new panel of anti-hOX40 mAb we generated a new knock-in (KI) mouse, designed to express hOX40 extracellular domain and mOX40 transmembrane and intracellular domains (Supplementary Fig. 1A). PCR confirmed integration of the construct and identified WT, hOX40KI^+/−^ and hOX40KI^+/+^ mice (Supplementary Fig. 1B). Analysis of OX40 surface expression (mouse and human) on resting splenocyte T cells from WT, hOX40KI^+/−^ and hOX40KI^+/+^ mice confirmed that the chimeric receptor was expressed at the cell surface on relevant cell types (Fig. 1A and B) and in a gene dose-dependent manner whereby hOX40KI^+/−^ expressed intermediate levels of both mouse and human OX40 and hOX40KI^+/+^ only expressed hOX40 (Supplementary Fig. 1C). Expression of OX40 in all three genotypes was largely restricted to T cell lineages (Fig. 1A, B and Supplementary Fig. 1D). Furthermore, in line with previous reports [14, 24–26], a hierarchal expression pattern amongst the T cell subsets was observed in all genotypes with expression highest on Tregs followed by CD4+ effectors, with limited expression on resting CD8+ T cells (Fig. 1A and B). To address whether the hOX40KI^+/+^ mice represented a functional model to study the OX40L:OX40 axis, we performed SPR analysis of mOX40L and hOX40L binding to hOX40 (Supplementary Fig. 1E). Both mOX40L and hOX40L bound similarly to hOX40, in agreement with earlier studies showing that mOX40L engages the same domains on hOX40 as hOX40L [27]. Furthermore, OX40 KO mice are reported to have a subtle defect in Treg numbers which is more apparent in young mice or under competition such as in bone marrow chimeras (reviewed in [28]). However expression of the chimeric receptor did not affect normal immune cell development (Supplementary Fig. 1F), further indicating that OX40L:OX40 signalling axis is intact in these hOX40KI mice.

**Figure 1.**
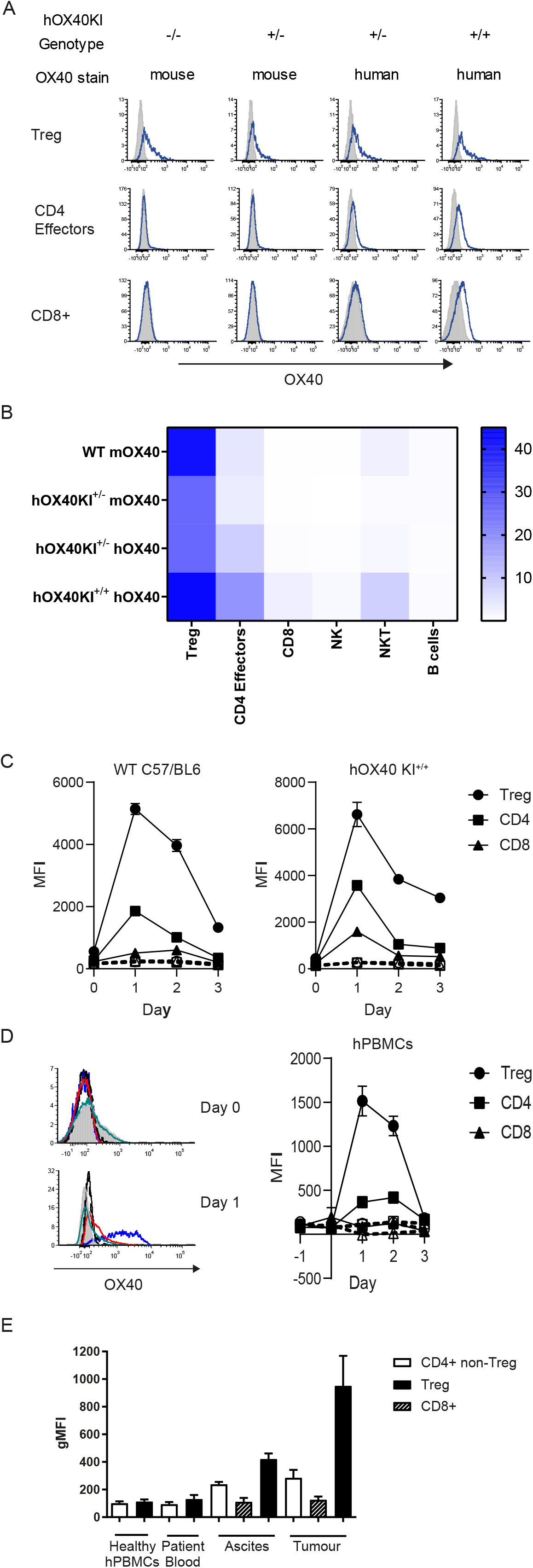
hOX40KI mice express hOX40 in a hierachial manner. A. Expression of OX40 (mouse (m) and human (h)) on Treg (top row), CD4+ Effectors (middle row) and CD8+ T cells (bottom row). Representative plots shown. B Heat map summarising OX40 expression as a percentage on resting mouse splenocytes (n=4). C. Expression of mOX40 (left panel) and hOX40 (right panel) on splenocytes from WT or hOX40KI mice activated with αCD3 and αCD28 (n=3). Isotype controls showed as dashed lines. D. Expression of hOX40 on hPBMCs activated with αCD3 and αCD28. Histograms show hOX40 expression on Tregs (blue line, isotype control black line), CD4+ (red line, isotype control black dashed line) and CD8+ (green line, isotype control grey filled histogram) from a representative donor on Day 0 (top panel) and Day 1 (bottom panel). Line graph (right panel) shows average expression, isotype controls shown as dashed lines (n=3). E. hOX40 expression on CD4+ effector T cells (white bars), CD8+ T cells (hatched bars) or Tregs (black bars) isolated from healthy donors or blood, ascites and tumour from cancer patients (n =4-16). Mean +/−SEM ****p<0.0001, ** p<0.01, * p<0.05 Tukey’s multiple comparison test.

Consistent with previous findings [29], activated splenocytes isolated from WT, hOX40KI^+/−^ and hOX40KI^+/+^ mice showed peak OX40 expression between 24-48 hours post activation (Fig. 1C). The kinetics of expression of hOX40 on hOX40KI^+/−^ and hOX40KI^+/+^ splenocytes correlated with that of hOX40 on activated human PBMCs (Fig. 1D). As with the resting cells, a hierarchy of expression was observed in both activated splenocytes and hPBMCs with greatest expression detectable on Tregs (Fig. 1C and 1D). Furthermore, samples taken from human cancer patients also showed the same pattern (Tregs > CD4+ effectors > CD8+ T cells) with highest OX40 expression on T cells isolated from tumour sites (Fig. 1E). Collectively, these results validated the use of the hOX40KI model as a tool to assess anti-hOX40 mAb.

### Generation and characterisation of a panel of anti-hOX40 mAb

A panel of seven anti-hOX40 mAb were subsequently generated by conventional hybridoma technology and further characterised. All mAb displayed a high affinity for hOX40 (KD values between 10^−9^-10^−10^M) as determined by SPR (Supplemental Fig. 2A) and did not bind mOX40 (Supplementary Fig 2B and C). We next sought to identify which domains of hOX40 these mAb bound. The crystal structure of hOX40 indicates four CRDs [27]. To determine which of these the anti-hOX40 mAb bound, hOX40 domain mutants were generated, lacking CRD1, CRD1+2 or CRD1, 2+3 (Fig. 2A). A hCD20 epitope tag and mOX40 CRD3 domain were added to the final construct to stabilise its expression. Across the panel of seven antibodies, at least one antibody bound to each of the different domains (Fig. 2B and Supplemental Fig. 2D). Further confirmation of specific binding domains was gained by performing cross-blocking experiments: anti-hOX40 mAb which bound to the same domain blocked the binding of one another e.g. both SAP 15-3 and SAP 28-3 bind CRD2 and blocked each other’s binding, whereas mAb binding to different CRDs could bind simultaneously (Fig. 2C). The domain binding for each antibody is summarised in Fig. 2D. Finally, the ability of the mAb to bind in the presence of the natural ligand, CD252 (OX40L) was investigated. The KD of hOX40:hOX40L is 0.25-1nM [30, 31] and the crystal structure of the complex demonstrates that the ligand spans CRD1-3 of OX40 [27]. SPR analysis revealed that only mAb binding to CRD4 were able to bind in the presence of the ligand (Fig. 2E and Supplemental Fig 2E), indicating ligand binding blocks binding of mAb that recognise epitopes in CRD1-3.

**Figure 2.**
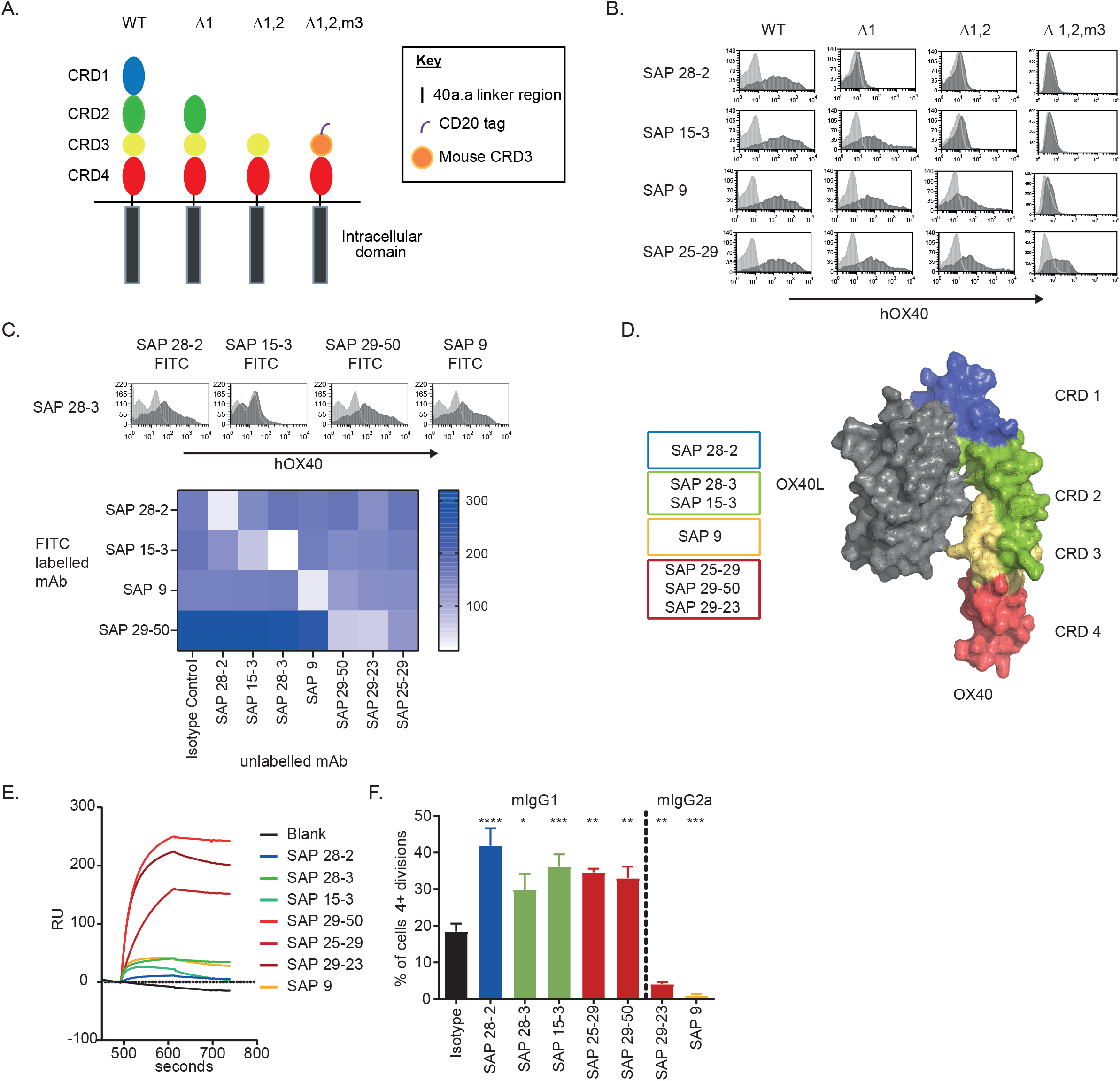
Characterisation of a panel of anti-hOX40 mAb. A. Schematic of the WT and domain mutant hOX40 constructs generated. CRD3 from mOX40 was used to stabilise the human CRD4 construct. B. Binding of anti-hOX40 mAb to domain constructs detected by a PE labelled secondary F(ab’)2 fragments. Representative histograms show hOX40 mAb binding (dark grey histogram) in comparison to an isotype control (light grey histogram). C. Representative histograms show anti-hOX40 FITC labelled mAb binding (dark grey histogram) in relation to an isotype control (light grey histogram) after binding of unlabelled anti-hOX40 mAb. The heatmap shows MFI of FITC labelled antibody binding in the presence of unlabelled antibodies with the absence of colour indicating blocking. D. Diagram summarising antibody binding domains in relation to the crystal structure of the OX40:OX40L complex. E. SPR analysis of anti-hOX40 mAb binding to hOX40-hFc in the presence of hOX40L-His fusion protein. F. Proliferation of hCD8+ T cells within PBMC cultures in response to sub-optimal anti-CD3 and anti-hOX40 mAb stimulation (representative of 4 individual donors). Mean +/− SEM ** p<0.01, * p<0.05 Dunnett’s multiple comparison test.

Antibodies were then assessed for their ability to augment sub-optimal anti-CD3 mediated proliferation of fresh hPBMC. All of the mIgG1 mAb increased proliferation of CD8+ T cells (Fig. 2F) whereas mIgG2a mAb reduced proliferation. Differences between mIgG1 and mIgG2a isotypes have been previously reported for other TNFR family members [32, 33] and so we class-switched the mAb so that both mIgG1 and mIgG2a isotypes were available.

### anti-hOX40 mIgG1 mAb are agonistic *in vivo*

We next sought to investigate the ability of the anti-hOX40 mAb to cause antigen-specific CD8^+^ T cell expansion *in vivo* making use of the OT-I model whereby antigen specific T cells are transferred into naive recipients. hOX40KI^+/−^ OT-I T cells, which have a TCR specific for OVA257–264/H-2K^b^ complex, were adoptively transferred into hOX40KI^+/+^ mice before vaccination with OVA and administration of anti-hOX40 mAb as either mIgG1 or mIgG2a (Fig. 3A). Both mIgG1 and mIgG2a anti-hOX40 mAb expanded antigen-specific CD8+ OT-I T cells in blood compared to OVA alone and to a similar extent with the exception of SAP 28-2 which was slightly weaker as a mIgG1 and significantly weaker as a mIgG2a (Fig. 3B and Supplementary Fig. 3A). However, despite reaching similar frequencies at the peak of the primary response, upon re-challenge with SIINFEKL peptide, a significantly smaller recall response was seen in mice that had received mIgG2a antibodies (Fig. 3B and Supplementary Fig. 3B). The frequency of OT-I cells pre-recall in mIgG1 treated mice strongly correlated with the strength of the recall peak (Supplementary Fig. 3C), with a similar relationship in mIgG2a treated mice. These data suggest that the number of OT-I cells present before re-challenge is a key determining factor for the strength of the recall response and that mIgG1 and mIgG2a mAb deliver signals during the primary response which results in different resting memory populations.

**Figure 3.**
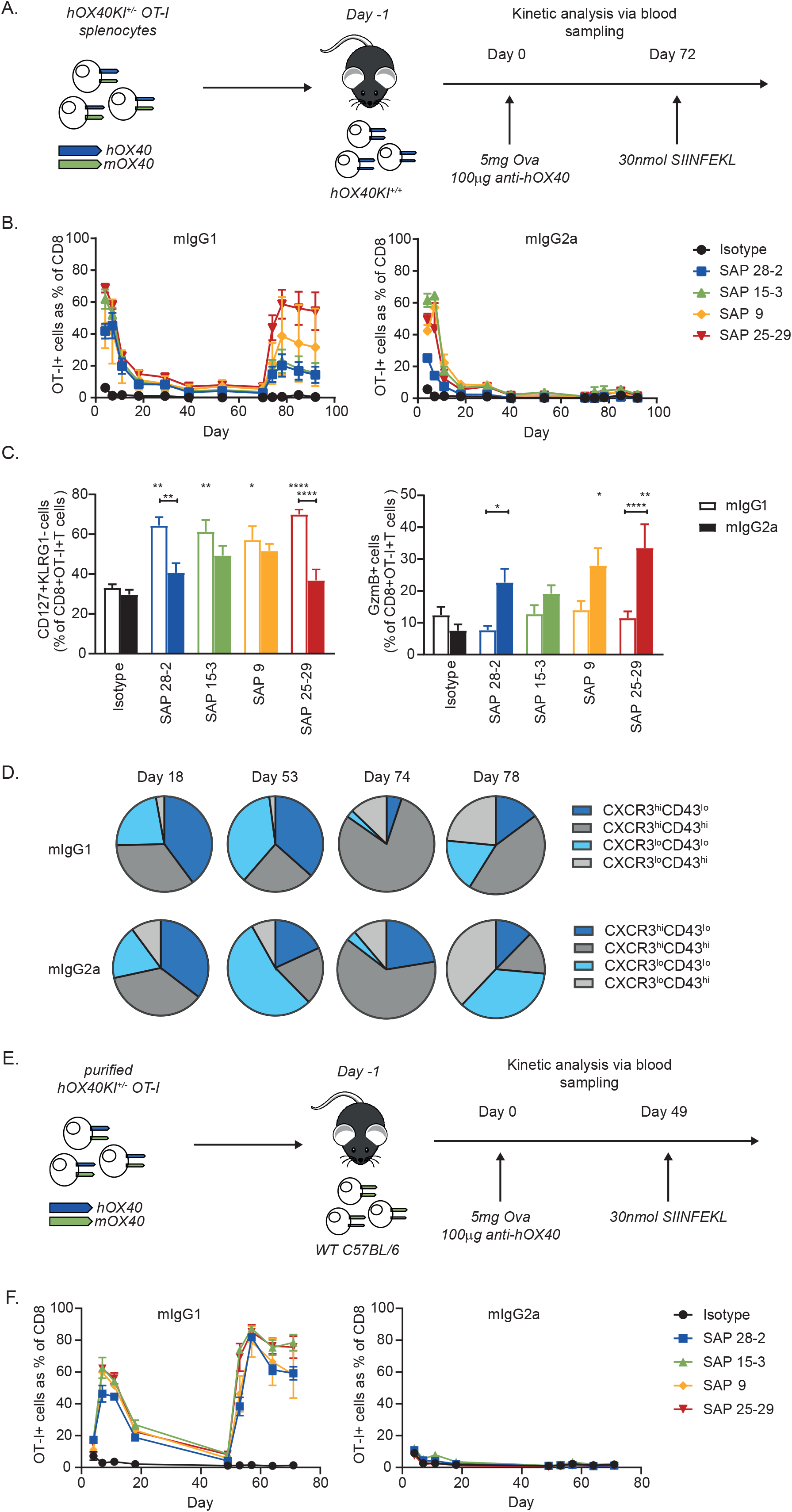
anti-hOX40 mIgG1 act agonistically in vivo. A. Schematic of the OT-I model used in B-D. 1×105 hOX40KI+/− OT-I cells were transferred into hOX40KI+/+ recipients. Mice were challenged with 5mg Ova and 100μg of antibody. B. Kinetic analysis of OT-I expansion in response to anti-hOX40 mIgG1 mAb (left panel) or mIgG2a mAb (right panel) (n=4 representative of 2 independent experiments). C Analysis of memory and effector phenotyping of OT-I+ T cells in blood at Day 18. MPECs - CD127+KLRG1-(left panel) and Granzyme B+ OT-I T cells (right panel) (n=4-8, pooled from 2 independent experiments). D. CXCR3/CD43 profiling of OT-I T cells in blood at timepoints indicated in response to SAP 25-29 stimulation (n=4). E. Schematic of the OT-I model used in F. 1×105 hOX40KI+/− OT-I were transferred into WT C57BL/6 recipients. Mice were challenged with 5mg Ova and 100μg of antibody. F. Kinetic analysis of OT-I expansion in response to anti-hOX40 mIgG1 mAb (left panel) or mIgG2a mAb (right panel) (n=4 representative of 2 independent experiments). Mean +/− SEM ****p<0.0001, *** p< 0.001, ** p<0.01 * p<0.05 Sidak’s multiple comparison test.

To understand this difference in memory response we phenotyped the OT-I T cells in the blood at both the primary and memory stages. We first assessed the ratio of short-lived effector cells (SLECs) to memory precursor effector cells (MPECs) as defined by the expression of CD127 and KLRG1 during the primary response [34]. As shown in Fig. 3C, the frequency of MPECs (CD127+KLRG1-) was higher in the mIgG1 groups at Day 18. Whilst frequencies of SLECs (CD127-KLRG1+) in the blood was similar between isotypes (Supplemental Fig 3D) granzyme B production was higher in SAP 9 and 25-29 mIgG2a treated mice compared to mIgG1 treated mice (Fig. 3C). These data also suggested that there may be a domain related trend in granzyme B production in those mice treated with mIgG2a isotypes, with the antibody which bound to CRD4 (SAP 25-29) producing more than those which bound to CRD3 (SAP 9) which in turn produced more than the antibody which bound to CRD2 (SAP 15-3) and CRD1 (SAP 28-2). In infection models, the relative frequencies of each subpopulation (SLECs vs MPECs) in the primary response does not always correlate with the accumulation of CD8+ cells during a recall response [34–36]. We therefore expanded our analysis to CXCR3 and CD43, shown in a lung infection model [37], to define three distinct populations of memory cells [38] with a hierarchy of recall response (CXCR3^hi^CD43^lo^ > CXCR3^hi^CD43^hi^ > CXCR3^lo^CD43^lo^) inversely correlated with activation markers such as KLRG1 [37]. In the memory phase of our OT-I model, mice that had been treated with mIgG1 anti-hOX40 mAb gave rise to a higher frequency of CXCR3^hi^CD43^lo^ and CXCR3^hi^CD43^hi^ (more proliferative) cells, in comparison to mice that had been treated with mIgG2a mAb (Fig. 3D and Supplemental Fig. 3E). This effect was most strongly seen with the SAP 25-29 CRD4-binding antibody (Fig. 3D and Supplemental Fig. 3E). This difference in frequencies between mIgG1 and mIgG2a treated mice whilst slight during the contraction phase (D18), became more evident during the resting memory period prior to re-challenge (D53). Immediately following re-challenge with SIINFEKL peptide (D74), both mIgG1 and mIgG2a groups displayed expansion of cells with high proliferative capacity but further into the memory response (D78) the hierarchy of mIgG1 treated mice having greater frequency of cells with greater proliferative capacity was re-established (Fig. 3D). Furthermore, mice that had been treated with mIgG2a anti-hOX40 mAb had a higher frequency of effector-like memory cells, denoted by CXCR3^lo^CD43^lo^ expression in the resting memory phase (D53) (Fig 3D). Again this contrast between mIgG1 and mIgG2a treated mice was re-established from D78 following re-challenge. These data indicate, that in our OT-I transfer model, the choice of isotype significantly impacts the development of a robust memory response.

These results also highlighted a discrepancy in the effect of the mIgG2a anti-hOX40 mAb between *in vitro* and *in vivo* experiments. mIgG2a anti-hOX40 mAb caused an inhibition of proliferation *in vitro* (Fig. 2F), whereas T cell expansion was seen *in vivo* (Fig. 3B). To probe the mechanisms of both mIgG1 and mIgG2a mAb to cause expansion of antigen-specific T cells, hOX40KI^+/−^ CD8+ OT-I T cells were purified and transferred into WT C57BL/6 mice (where they represent the only cell type expressing hOX40) to see if they were both capable of acting directly on CD8+ T cells (Fig 3E). Whilst mIgG1 mAb were able to drive a similar expansion as before, mIgG2a mAb had no effect (Fig. 3F). These results indicate that the mIgG2a mAb act indirectly, through non-CD8+ T cells, to facilitate OT-I expansion in contrast to the ability of mIgG1 to act directly on the CD8+ OT-I T cells.

### anti-hOX40 mIgG2a mAb deplete hOX40 expressing cells

We hypothesised that the indirect effect of mIgG2a mAb might involve deletion of Treg via activatory FcγR-mediated ADCC and ADCP effector functions [32]. To determine if the anti-hOX40 mIgG2a mAb were evoking Treg deletion, spleens were harvested from hOX40KI^+/+^ mice four days after treatment with either anti-hOX40 mIgG1 or mIgG2a mAb (Fig. 4A). The number of cells present in different T cell subsets in the spleen was assessed and it was clear that the mIgG2a mAb were able to significantly reduce the number of Tregs and to a lesser extent CD4+ effectors whereas mIgG1 treated mice evoked expansion of T cell populations (CD8+ and OT-I) (Fig. 4B and Supplemental Fig. 4A) with the greatest expansion seen in the OT-I population. The relative deletion of T cell populations in the mIgG2a treated mice correlated with the amount of hOX40 surface expression seen following activation *in vitro* (Fig. 1C) as the most significant difference was seen in the Treg population followed by CD4+ effectors and lastly CD8+ cells. In contrast, the number of CD8+ and OT-I cells had the largest increase in proportion following treatment with the mIgG1 mAb. The CD8:Treg ratio was unchanged following treatment with the mIgG1 mAb. However SAP 25-29 mIgG2a produced a significant increase in the CD8:Treg ratio with a trend towards an increase also seen for SAP 15-3 mIgG2a and SAP 9 mIgG2a (Fig. 4B). Interestingly SAP 9 mIgG2a also showed a decrease in inter-sample variability when looking at the number of OT-I cells and a trend towards a reduction in CD8+ T cell numbers, indicating superior deletion capacity compared to other mAb. Therefore, we next explored the ability of SAP 9 to delete human Tregs *in vivo*. Un-activated human Treg do not typically express appreciable OX40 levels (Fig. 1D, [39] and reviewed in [29]) and so to upregulate expression we performed a NOG:PBMC transfer model in which hPBMC are first activated (through a xenoresponsive graft versus host response) before being transferred into a new NSG recipient mouse where deletion can be assessed. The transferred Treg upregulate hOX40 to levels similar to those observed within tumours and significantly above the levels of CD8+ T cells (Supplemental Fig. 4B). Mice harbouring the NOD phenotype exhibit reduced NK cell frequency and function and the absence of hemolytic complement activity [40], leaving myeloid FcγR bearing cells and ADCP as the principal mAb effector mechanism. Using this approach, SAP 9 hIgG1 was shown to be capable of deleting human Tregs at least as well as the clinically-approved Treg deleting anti-CTLA-4 mAb Yervoy. (Fig. 4D); thereby augmenting the CD8:Treg ratio (Fig. 4E). Furthermore, when using a humanised version of SAP 9 hIgG1, significant depletion of human Tregs was observed (Fig. 4F), alongside a significant improvement in the CD8:Treg ratio (Fig. 4G), unlike with CAMPATH-1 which deleted all T cells. These data indicate that with the correct isotype and significant expression of hOX40, anti-hOX40 antibodies are capable of specifically depleting Tregs and improving CD8:Treg ratios *in vivo*.

**Figure 4.**
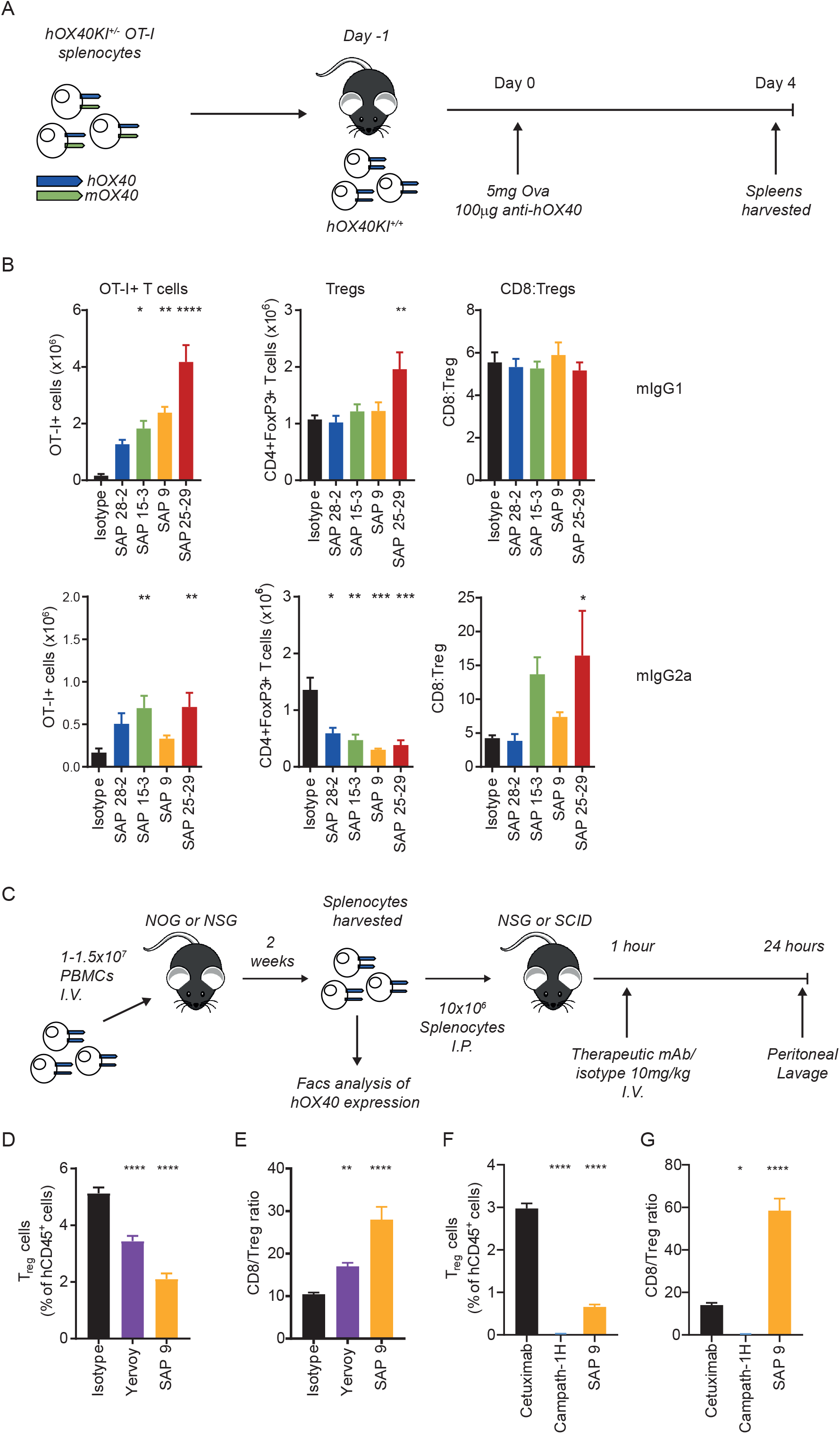
anti-hOX40 mIgG2a mAb elicit Treg cell depletion in vivo. A. Schematic of OT-I model used to assess cell depletion. B. Assesment of Tetramer+ve (left panel), Treg cell numbers (middle panel) and CD8:Treg ratio (right panel) in response to anti-hOX40mAb either as a mIgG1 (top row) or mIgG2a (bottom row) (n=4-6 pooled from two independent experiments). C. Schematic of the NSG/PBMC model to assess depletion of hPBMCs. 1.5 ×107 (D, E) or 1 ×107 (F, G) hPBMC were transferred into NOG (D, E) or NSG (F, G) mice. 2 weeks later splenocytes were harvested and transferred into SCID (D,E; n=11-13 pooled from 2 independent experiments) or NSG (F, G; n=7 pooled from 2 independent experiments) recipients which were then treated with depleting antibodies. Treg numbers (D, F) and CD8:Treg ratio (E, G) were determined by flow cytometry. Mean +/− SEM, ****p<0.0001, *** p< 0.001, ** p<0.01, * p<0.05 Sidak’s (B) and Tukey’s (F and G) multiple comparison test.

### Extent of mIgG1 agonism and mIgG2a deletion correlates with OX40 domain specificity

Throughout the data detailed above it became apparent that the strength of deletion/agonism was associated with the domain of hOX40 bound by the various mAb. To assess this further we grouped our results into membrane distal (CRD1+2) and membrane proximal (CRD3+4) binding mAb (Fig. 5A and B). We also assessed groups reflecting those able (CRD4) or unable to bind in the presence of ligand (CRD1+2+3) to see if there was a correlation with ligand competition (Fig. 5C and D). The strength of agonism seen with the mIgG1 mAb was highest for membrane-proximal binding mAb for both expansion of CD8+ OT-I cells and Tregs (Fig. 5A). In contrast, the expansion of CD8+ OT-I cells with the mIgG2a isotype did not correlate with domain, although the depletion of Tregs did, being highest for membrane proximal domains (Fig. 5B). Likewise, mAb binding outside of the ligand-binding domain displayed the highest level of mIgG1-mediated agonism (ability to expand both CD8+ OT-I cells and Tregs) (Fig. 5C). Interestingly the ability of mIgG2a mAb to to deplete Tregs did not significantly correlate with binding to CRD4 (Fig 5D), i.e. outside of the OX40L binding region.

**Figure 5.**
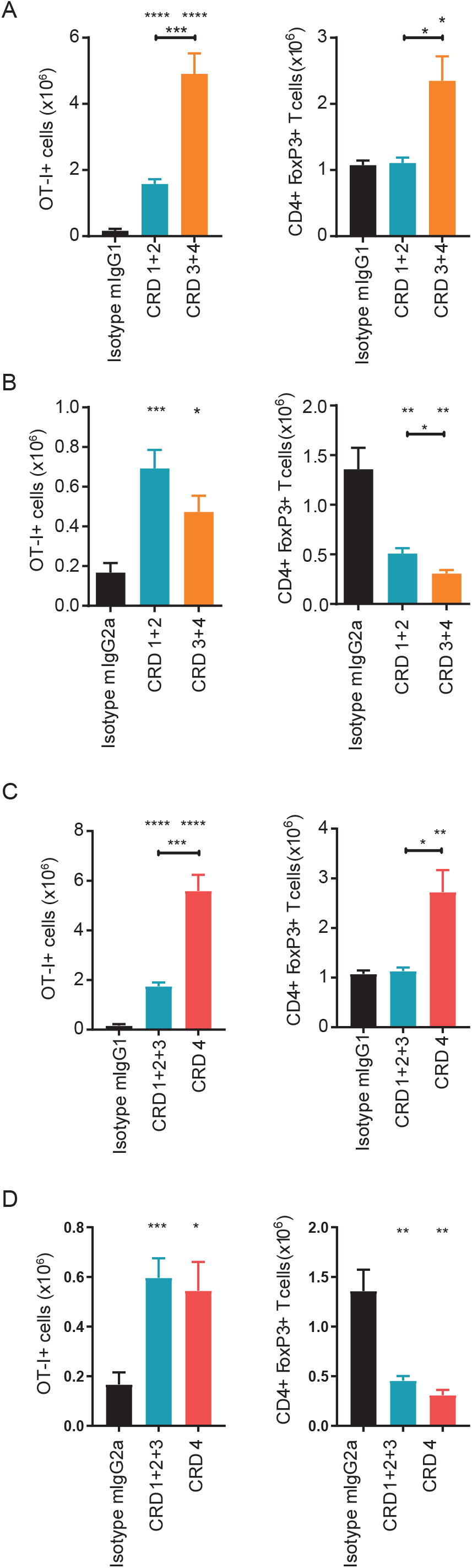
Anti-hOX40 mAb agonism and depleting activity are related to domain binding. Model used as in Fig 4A and B. A. Analysis of OT-I cell numbers (left panels) and Treg cell numbers (right panels) in response to anti-hOX40 mIgG1 mAb grouped into proximal binders – CRD3+4 or distal binders – CRD1+2. B. As in A except mIgG2a mAb was used. C. Analysis of OT-I cell numbers (left panels) and Treg cell numbers (right panels) in response to anti-hOX40 mIgG1 mAb grouped into those which can bind in the presence of ligand - CRD4 vs those which cannot - CRD1-3. D. As in C except mIgG2a mAb was used. Mean +/− SEM ****p<0.0001, *** p< 0.001, ** p<0.01, * p<0.05 Tukey’s multiple comparison test (A and B; n=6-16) (C and D; n=6-19).

### Ability of hOX40 mAb to control tumour growth

We next sought to determine the immunotherapeutic potential of the anti-hOX40 mAb *in vivo* in our hOX40^+/+^ KI mice, evaluating a mAb which bound to each CRD of hOX40 as both a mIgG1 and mIgG2a isotype. hOX40^+/+^ KI mice were inoculated with OVA expressing E.G7 lymphoma cells and subsequently treated with anti-hOX40 mAb once tumours had established. anti-hOX40 mAb, as both mIgG1 and mIgG2a, were able to elicit tumour control, with mice eradicating established tumours in most treatment groups (Fig. 6A). Nonetheless, with the exception of SAP 25-29, the mIgG1 mAb caused a higher % survival than the mIgG2a mAb, although there was no obvious domain preference amongst the antibodies in terms of anti-tumour activity. Importantly, mice also appeared to form durable memory responses as upon rechallenge no mice developed a secondary tumour (Supplemental Fig 5A).

**Figure 6.**
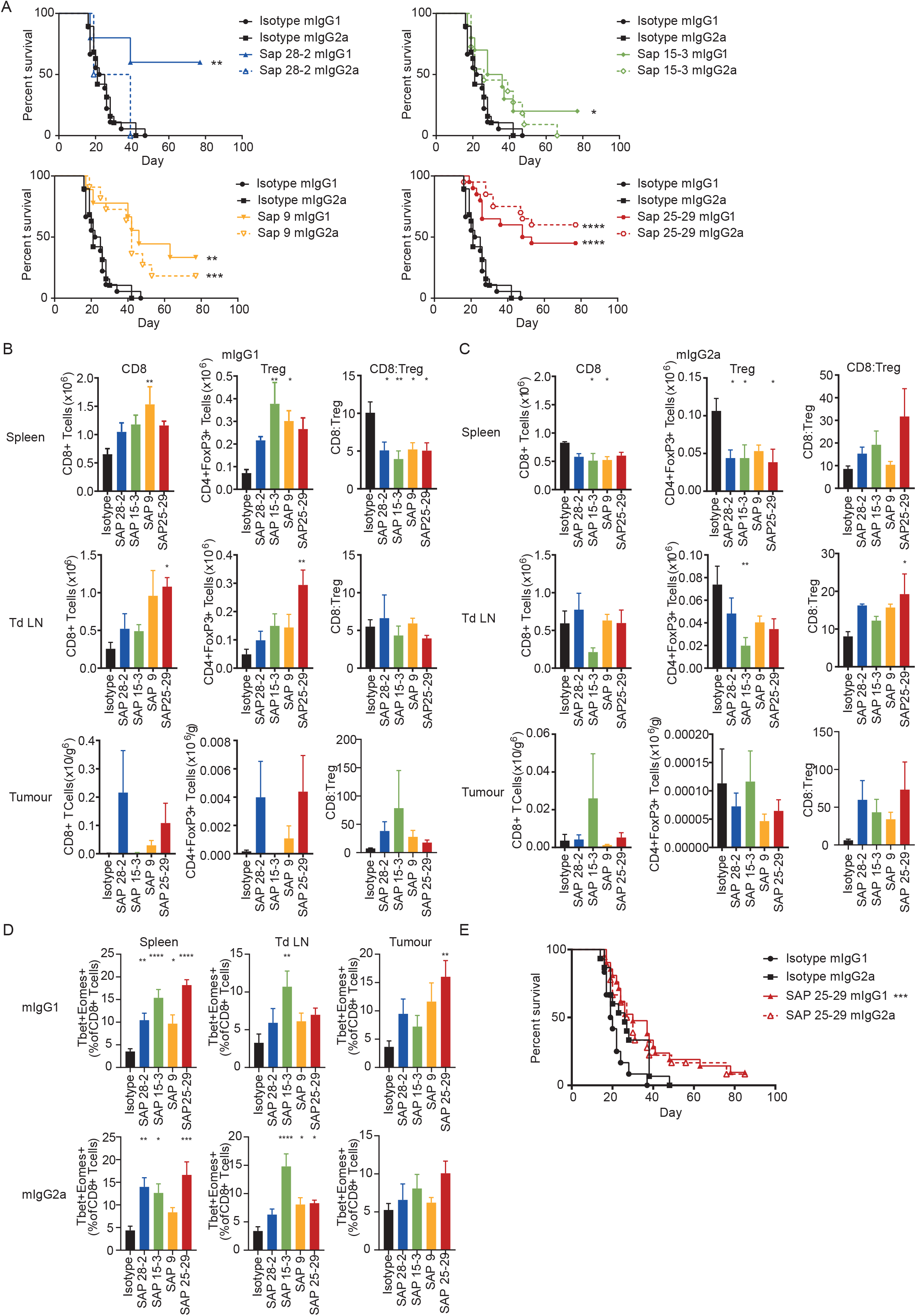
anti-hOX40 mAb are therapeutic as both mIgG1 and mIgG2a. A. Survival curves for mice challenged with E.G7 lymphoma cells (0.5×106) and treated with 3×100μg mAb once tumours are between 5×5 and 10×10mm. Data pooled from 2 independent experiments (n=5-19). Assessment of T cell populations in spleen (top panels), Tumour draining lymph node (middle panels), and Tumour (bottom panels) isolated 24 hours post second mAb dose either as a mIgG1 (B) or mIgG2a (C). n=3-5, representative of 2 independent experiments. D. Analysis of Tbet and Eomes expression in CD8+T cells isolated from Spleen (left panels), Tumour draining lymph node (middle panels) and Tumour (right panels) from mice treated with either mIgG1 (top row) or mIgG2a (bottom row). Data pooled from 2 independent experiments (n=8-9). E. Survival graphs for mice challenged with MCA-205 cells (0.5×106) and treated with 3×100μg mAb once tumours were 5×5mm. Data pooled from 3 independent experiments (n=12-19). Mean +/− SEM ****p<0.0001, *** p< 0.001, ** p<0.01, * p<0.05 Log-rank (Mantel-Cox) for survival graphs A and E and Dunnett’s multiple comparison test for B-D.

To determine the mechanism of tumour control, mice were again inoculated with E.G7 lymphoma cells and treated with anti-hOX40 mAb before organs were harvested and assessed for changes in immune inflitrate. Consistent with data from the OT-I model, the general trend seen in mIgG1 treated mice was expansion of T cells (CD4+ effectors, CD8+, Tregs and OT-I T cells) whilst mIgG2a mAb caused T cell depletion (Fig. 6B and C and Supplemental Fig. 5B and C). This effect was most prominent in the spleen but also observed within the tdLN (CD4+ and Tregs). Cell numbers recovered from tumours were very small therefore it was difficult to ascertain clear trends within the TILs (Fig. 6B and C). Of particular importance was the CD8:Treg ratio as it is known that a high CD8:Treg within human tumours is associated with prolonged survival [41] and depletion of Tregs allows for anti-tumour immunity and rejection [42, 43]. Our results show that mice treated with mIgG2a anti-hOX40 tended to increase the CD8:Treg ratio within the spleen, tdLN, and tumour (Fig. 6C). Interestingly, although in the spleen the mIgG1 treated mice showed a significant reduction in the CD8:Treg ratio, in the tumour an increased trend was observed (Fig. 6B), it is unclear whether this discrepancy between locations occurs as a result of different response kinetics, with priming occurring in lymphoid organs and subsequent recruitment to the tumour or perhaps direct priming within the TME whereby the greater antigen density leads to a more rapid response and hence increase in CD8:Treg. A domain trend could also be seen within some of the T cell subsets; CD8+ OT-I depletion in the spleen and CD4+/Treg expansion within the tdLN associated with membrane proximal domains (Fig. 6B and C and Supplemental Fig. 5B and C).

To better understand these responses, next, we examined the levels of expression of the T-box transcription factors T-bet and eomesodermin (Eomes) which play roles in CD8+ T cell differentiation and function [44]. Both transcription factors cooperate to promote cytotoxic lymphocyte formation, which correlates with the upregulation of perforin and granzyme B in antigen specific cells [34, 45, 46], as well as sustaining memory phenotypes [47]. Examination of the percentage expression of T-bet and Eomes on T cells within the spleen, tdLN and tumour of anti-hOX40 treated mice revealed that T-bet and Eomes double positive cells increased in the CD8+ populations when mice were treated with either mIgG1 or mIgG2a anti-hOX40 mAb with very little expression detectable on the CD4+ populations (Fig. 6D and data not shown). A domain trend was also broadly observed in mIgG1 treated mice, most obviously within the spleen and tumour. The percentage of cells expressing the effector molecule, granzyme B, was largely unchanged on CD4+ T cells (data not shown). However CD8+ T cells both within the spleen and tumour saw an increase in granzyme B producing cells in both mIgG1 and mIgG2a treated mice in comparison to the isotype control (Supplementary Fig. 5D). These results show that all anti-hOX40 mAb, irrespective of isotype or domain binding region, are able to produce functional effector cells, as well as cells expressing transcription factors important for effector and memory cell formation within a tumour environment.

To determine if the ability of these mAb to evoke tumour control was consistent across tumour models we utilised the subcutaneous MCA-205 sarcoma. Using SAP 25-29 as a paradigm, we again saw both isotypes providing tumour control, albeit more limited in comparison to the E.G7 tumour model and with no isotype preference (Figure 6E).

## Discussion

In the current study, we generated and characterised a panel of mAb for the targeting of hOX40. They were shown to bind throughout the four different hOX40 CRDs and were expressed as both mIgG1 and mIgG2a isotypes to understand the effects of both isotype and domain binding on agonistic and therapeutic potential in a newly developed hOX40 KI mouse model.

The hOX40 KI mouse model displayed an expression pattern of hOX40 largely reflecting that seen on healthy hPBMCs and samples from ovarian cancer patients with a hierarchy of expression of Treg>CD4+>CD8+. Nevertheless, differences were observed such that higher levels of hOX40 were seen on peripheral CD8+ T cells in the homozygous KI mouse than would be expected on hPBMC. Expression on all cell types was gene-dose dependent being ~ 50% lower in heterozygous KI mice and so allowed assessment of expression level for any biological effects observed. In addition, constitutive expression of hOX40 was observed on peripheral Tregs in the mice, in contrast to negligible levels on resting hPBMCs. However, this expression pattern reflects that observed on Treg in TILS isolated from cancer patients and so appears a reasonable model for studies in oncology.

The panel of anti-hOX40 mAb collectively bound to all four CRDs of hOX40 but only mAb binding to the most membrane proximal domain (CRD4), were able to bind in the presence of the natural ligand OX40L (CD252), raising the possibility that certain effector functions could be influenced by concurrent ligand binding for the CRD4 binding mAb. In the hOX40 KI mouse, mOX40L not hOX40L is present which may have led to immune defects due to the absence of OX40L:OX40 interaction.

However, no overt differences in immune development or homeostasis were observed in the homozygous OX40 KI mice (Supplementary Fig. 1). Compann and Hymowitz demonstrated that mOX40L binds the same domains and makes similar contacts to hOX40 as hOX40L, despite their relatively low sequence homology (~40%) [27], indicating that the murine OX40L binds the human receptor and maintains OX40L:OX40 signals. It also indicates that the model is suitable to address any influences of ligand binding on the activity of the mAb panel. Although not studied directly, no overt effects appeared to be driven by the presence or absence of OX40L binding; i.e. all antibodies regardless of ligand blocking were able to elicit function in an OT-I transfer model and in a number of tumour models.

It is well known that isotype helps dictate mAb effector function due in part to differences in FcγR interactions, and so isotype choice is important for delivering therapeutic efficacy according to the mAb mechanism of action [32, 48, 49]. For example, the mIgG2a isotype is optimal for direct targeting antibodies, which rely on efficient activatory FcγR interaction and effector stimulation for target cell deletion, whereas immunostimulatory mAb are often optimal as mIgG1, which preferentially engage the inhibitory FcγRII to facilitate receptor cross-linking and agonistic signalling in the immune cell [32, 50]. These paradigms have also held true for TNFR family members; mIgG1 antibodies being agonistic via engagement of FcγRII, with mIgG2a either inhibitory or with limited agonistic effects yet capable of activatory FcγR-mediated target cell depletion [14, 24, 32, 33]. Our initial results revealed a fundamental difference in effect between mIgG1 and mIgG2a anti-hOX40 mAb; in hPBMC proliferation assays mIgG2a mAb resulted in a decrease in the percentage of proliferating T cells compared to the isotype control whereas mIgG1 counterparts evoked increases in T cell proliferation. These disparate responses were not reflected in our hOX40^+/−^ OT-I transfer studies in hOX40^+/+^ KI mice, where both mIgG1 and mIgG2a anti-hOX40 mAb were able to cause strong expansion of antigen-specific CD8+ OT-I cells regardless of domain binding location. However, subsequent studies revealed that mIgG2a mAb activity was dependent upon transfer into hOX40KI^+/+^ mice. When hOX40^+/−^ OT-I cells were transferred into WT mice the mIgG2a mAb were no longer able to elicit expansion of CD8+ OT-I cells, unlike the mIgG1 mAb, which retained this activity. Together, these results indicate that the mIgG1 mAb cause direct agonism on the hOX40^+/−^ OT-I cells resulting in their expansion, whereas the mIgG2a mAb require hOX40 expressing non-CD8+ cells to provide expansion.

Further to this, and in agreement with our initial studies on hPBMC, enumerating the subpopulations of T cells within the spleen of hOX40^+/+^ KI mice revealed that the mIgG2a anti-hOX40 mAb uniquely caused depletion of Tregs. Lower levels of depletion were seen in the CD4+ effector population with little-to-no effect in the general CD8+ population, despite the expansion of antigen specific CD8+ OT-I cells, as seen in the blood. In contrast, the mIgG1 anti-hOX40 mAb caused expansion of all T cell populations, most significantly in the general CD8+ and OT-I populations, with relatively less expansion in the Treg and CD4+ populations. Assuming the presence of relevant effector cells and normal distribution of FcγRs in our OT-I model, it is likely that differential FcγR interactions of the different isotypes explains their disparate effects. Furthermore the superior affinity for the activatory FcγR and resultant deletion of suppressive Treg cells seen in mice treated with mIgG2a mAb likely explain the mechanism behind the expansion of CD8+ OT-I cells in the blood and spleen of hOX40^+/+^ KI but not WT mice.

Another disparity seen between the anti-hOX40 mIgG1 and mIgG2a treated mice was the percentage of memory cells generated after hOX40^+/−^ OT-I cells were transferred into hOX40KI^+/+^ recipients and rechallenged with SIINFEKL peptide. Considering the similar levels of CD8+ OT-I cells at the peak of the primary response it was surprising to observe such a difference in the frequency of CD8+ OT-I cells in the memory phase. A positive correlation between the frequency of CD8+ OT-I cells pre-recall and the frequency of CD8+ OT-I cells at the peak of the memory response highlighted the possibility that it was simply the result of the amount of cells present at the time of re-challenge, with mIgG1 mAb but not mIgG2a mAb providing signals for survival/persistence. To explore this further, we phenotyped the CD8+ OT-I cells within the primary, resting memory, and memory phases of the response. We found that within the primary response the MPEC population as described by CD127^+^KLRG1^−^ expression was increased at day 18 in mice treated with mIgG1 mAb compared to those treated with mIgG2a. Mice treated with mIgG1 mAb also generated a higher percentage of CXCR3^hi^CD43^lo^ highly proliferative cells in comparison to mice treated with the mIgG2a mAb. These findings, coupled with those described above relating to the frequency of antigen specific cells pre-rechallenge explain the disparity in the memory response comparing mIgG1 versus mIgG2a mAb. However, an alternative, but not mutually exclusive explanation could involve the higher percentage of SLECs and granzyme B producing cells observed in the primary response of mIgG2a treated mice, as well as the higher frequency of effector-like (CXCR3^lo^CD43^lo^) memory cells. The exact mechanisms underpinning this dichotomy are not immediately clear but one possibility is that mIgG2a deplete Tregs alongside, to a lesser extent, CD4+ effectors, to influence priming. Importantly though this higher proportion of potent effector and effector-like cells in mIgG2a treated mice versus highly proliferative cells in mIgG1 treated mice may together explain these differences in memory and moreover provide a rationale for how both isotypes are able to cause equivalent efficacy in mouse tumour models, despite exhibiting opposing mechanisms of action.

In addition to the isotype dependent effector functions discussed above, domain binding of the mAb revealed a significant effect on the strength of both mIgG2a depletion and mIgG1 agonism. anti-hOX40 mAb which bound to CRD4 were more potent agonists as mIgG1 when compared to mAb which bound to CRD1-3. This result is in direct contrast to our previous observations with anti-CD40 mAb [22] where CRD1 binding mAb were more agonistic. However, for both the anti-hCD40 and anti-hOX40 mAb, optimal agonistic function correlated with binding outside the natural ligand binding region. This potentially suggests that the combined effects of both the ligand and the mAb in clustering the receptor are required to elicit optimal agonism. This conjecture is partially supported by recent evidence from Zhang et al. who also demonstrated strong agonistic function with a CRD4 binding anti-mOX40 mAb. However, those authors also documented equivalent agonistic activity with a CRD2-binding, ligand blocking anti-mOX40 mAb [23], indicating the fine epitope is also important, as we reported previously for anti-CD40 [22].

In comparison to delivering agonism as mIgG1, mIgG2a anti-hOX40 mAb evoked potent deletion of Treg in the hOX40 KI mice. The ability of TNFR targeting mAb to cause depletion of intratumoral Tregs has been clearly demonstrated in recent years [10, 14–16], both increasing their therapeutic possibilities but confusing potential mechanisms of action. To allow us to assess, in a more translational setting, the depleting ability of our anti-hOX40 mAb, we performed depletion experiments in NSG mice engrafted with human target cells expanded from PBMCs. In these experiments, hOX40 and other TNFR family members become upregulated on the activated Treg as they mount xeno-reactive responses (Supplemental Fig. 4B and [17] - enabling them to serve as targets akin to those seen in tumour samples. Specifically, we assessed the ability of SAP 9 hIgG1 to delete Treg and compared it against the clinically relevant Treg depletor, the anti-hCTLA-4 mAb, Yervoy [51]. SAP 9 was not only as strong a Treg depleter but also generated a higher CD8:Treg ratio in comparison to Yervoy indicating the therapeutic potential of this anti-hOX40 mAb. Importantly, deletion of Treg was specific even though all T cell subsets had expanded and were activated through the NSG passage.

Having established the agonistic and depletory capacity of our panel of mAb, we next explored them in two different solid tumour models, showing that the therapeutic effects of both mIgG1 and mIgG2a mAb were similar, with no clear benefit attributed to domain binding.

The ability of both isotypes to act therapeutically is perhaps surprising but likely reflects their relative propensity to elicit different effector functions in the tumour microenvironment (TME). The latter is known to have profound effects on therapeutic efficacy [52]. For example, if a tumour has a high infiltrate of cells expressing activatory FcγR such as NK cells and macrophages, mIgG2a mAb and depletion of detrimental target cells may be favoured over the T cell-agonising activity of a mIgG1 mAb. Conversely, if the inhibitory FcγRIIB is more prevalent within the cellular infiltrate, mIgG1-mediated agonism may be more prominent. Lymphoid organs outside the tumour may also be relevant. In the E.G7-Ova model, although the magnitude of effect differed, the mIgG1 mAb largely caused T cell expansion and the mIgG2a depletion in both the spleen and draining lymph node. It seems likely therefore that these intrinsic differences underpin the relevant mechanism of action in each case. Accordingly, mIgG2a reagents likely achieve therapeutic effects through depleting Tregs, releasing T cell effector responses whereas mIgG1 expand T cell numbers and hence increasing cytotoxic effectors within the tumour site.

One additional point of difference between the mIgG1 and mIgG2a reagents was the induction of antigen-specific memory: In the E.G7-Ova lymphoma model both mIgG1 and mIgG2a isotypes were able to elicit tumour control and generate memory sufficient to prevent tumour growth upon rechallenge. This suggests that despite the marked difference in the memory response seen in the OT-I model (mIgG1>>mIgG2a), other factors must be operational in the presence of tumour to account for the ability of both mAb isotypes to generate protective memory responses. One possibility is that the threshold for memory is relatively low in the presence of an immunogeneic tumour and that the limited level of memory recall seen following treatment with the mIgG2a is sufficient. Alternatively, it may indicate that the mIgG2a mechanism of action, deleting Tregs, allows the expansion of otherwise cryptic epitopes allowing T cell control as was previously described in the CT26 model [53].

In summary, our findings show that immunomodulatory mAb directed to hOX40 can harness multiple mechanisms of action to elicit tumour control. They also show that these mechanisms can be modulated dependent upon the choice of isotype and domain binding region. Targeting the membrane proximal domains appears optimal for both deletion and agonism; with the latter strongly driven by isotypes with low A:I ratio such as mIgG1. Lowering the A:I ratio can be attained in many different ways, for example by increasing the affinity for FcγRIIB, which has been shown to mediate more effective agonism for anti-CD40 mAb [54] but as yet it is unclear whether this will be true *in vivo* for anti-hOX40 mAb. These findings have implications for the design of the next generation of anti-hOX40 mAb for the clinic. Current reagents typically display an unmodified hIgG1 isotype and although safe have not delivered strong anti-tumour effects [7]. These hIgG1 reagents would be expected to deliver the Treg deleting function indicated here but to date this activity has not been shown clearly in patients, with the topic currently debated for other Treg targets such as CTLA-4 [55–57]. Furthermore, their deletion may not be sufficient to elicit tumour regression in most human cancers, unlike the mouse models shown here. Therefore, mAb with appropriate isotypes and further engineering (hIgG2B[22, 58], SELF[54, 59], V11[54, 60]) to elicit potent T cell agonism may be warranted for further investigation.

## Methods and Materials

### Mice

C57BL/6 mice and OT-I transgenic mice were obtained from Charles River Laboratories. NSG mice were purchased from Jackson Laboratories. hOX40 knock-in mice were generated by Ozgene. hOX40/OT-I mice were generated by crossing homozygote hOX40 KI with homozygote OT-I mice. PBMC-NOG/SCID mice (primary human xenograft model) were generated by injecting NOG or NSG mice with 1-1.5×10^7^ PBMC isolated using Ficoll-Paque PLUS in 200μl PBS. SCID or NSG mice were subsequently injected with 10×10^6^ splenocytes from reconstituted NOG or NSG mice. For all experiments young adult mice were sex- and age-matched and randomly assigned to experimental groups. All procedures were conducted in accordance with UK Home Office guidelines and were approved by the University of Southampton’s ethical committee or at BioInvent under Dnr 14760/2016.

PCR genotyping of mice hOX40 KI was detected using the following primers F – AGTGCCCACGCTTCCTGAGGA and R - CTTGAGGATGCCAGAGGAGGC which give a 290bp product. mOX40 was detected using the following primers F - TCTCCACCCACCTTGGTGACT and R – GCCAGCAGGACAGTCAAGGA which give a 174bp product.

### Human Samples

Peripheral blood mononuclear cells (PBMCs) were obtained from healthy adult volunteers from either Southampton National Blood Service, UK or Hallands Hospital Halmstad, Sweden. For clinical samples, ethical approval was obtained from the Ethics Committee of Skåne University Hospital, Sweden. For NSG reconstitution experiments done in Southampton, hPBMCs were purchased from STEMCELL Technologies.

### Antibody Production and labelling

Anti-hOX40 mAb were created using conventional hybridoma technology [61]. Isotype switching was carried out by subcloning variable regions into expression vectors containing constant regions of different antibody isotypes as previously described [62]. Constructs were transfected into mammalian cell lines and mAb purified either using a protein-G or a protein-A column. All preparations were deemed endotoxin low (<1ng/mg protein) before use. FITC-labelling: FITC (Sigma) was dissolved in bicarbonate buffer at a concentration of 2mg/ml. Anti-hOX40 mAb were FITC labelled at a ratio of 1:10 FITC:mAb. The FITC labelled mAb were purified using a desalting column (GE Healthcare). Cetuximab was a kind gift from Thomas Valerius and Campath-1 was a kind gift from Geoff Hale.

### Humanisation of SAP 9

The variable regions of SAP 9 heavy and light chains were sequenced from the hybridoma by PCR. The sequence was humanised using Macromoltek’s proprietary humanisation algorithms. A generic antibody signal peptide sequence was then added to the humanised variable region sequences and the amino acid sequences converted into nucleotide sequences using https://www.bioinformatics.org/sms2/rev_trans.html. Nucleotide sequences were synthesized by GeneArt and subcloned into expression vector pEE6.4 (Lonza) for expression.

### Binding domain determination/ blocking experiments

hOX40 constructs were transiently transfected into 293F cells before addition of 10μg/ml anti-hOX40 mAb. For domain binding determination PE-labelled secondary anti-mouse Fc antibody (Jackson Laboratories) was added. For blocking experiments unlabelled antibody was added for 30 minutes before addition of a second FITC-labelled anti-hOX40 mAb. Binding of PE- or FITC-labelled antibodies was assessed using flow cytometry (see below).

### Surface Plasmon Resonance

A Biacore T100 upgraded to a T200 (GE Life Sciences) was used to measure hOX40:mAb and hOX40:hOX40L:mAb interactions. 1μg/ml of hOX40-hFc was immobilised onto a CM5 chip (GE Healthcare) and a range of concentrations from 0-500nM of analyte (anti-hOX40 mAb) was injected over the chip. The chip was regenerated between each run using 10mM glycine pH 1.5. For ligand blocking experiments 100nM of hOX40L-His was immobilised onto a CM5 chip coated with an anti-His mAb. 100nM hOX40-hFc was subsequently captured followed by the injection of anti-hOX40 mAb (15μg/ml).

### *In vitro* assays

Human PBMCs were isolated using lymphoprep separation and used either fresh or frozen prior to use. Cells were frozen in 90%FCS, 10%DMSO at 5×10^7^ in 1.5ml, then stored in liquid nitrogen. For human proliferation assays using fresh cells; cells were labelled with 1μM CFSE immediately following isolation and prior to culture at high density (1.5×10^7^ cells/ml) for 2 days at 37°C prior. Cells plated at 1×10^5^ cells/well in a 96 round bottomed well plate and stimulated with soluble anti-CD3 (OKT3, 5ng/ml) and anti-hOX40 mAb (5μg/ml). Cells were stained with appropriate antibodies and CFSE dilution assessed via flow cytometry. For expression assays frozen hPBMCs were thawed and rested overnight and then stimulated with plate bound anti-CD3 (OKT3, 15ng/ml) and anti-CD28 (CD28.2, 0.5μg/ml). Cells were harvested, stained with appropriate antibodies and assessed via flow cytometry.

For mouse expression assays, splenocytes were isolated using a 100μm strainer, washed in PBS and red cell lysed using ACK lysis buffer. Cells were re-suspended in complete RPMI (FCS (10%), Glutamine (2mM), Pyruvate (1mM), Penicillin (100units/ml), Streptomycin (0.1mg/ml) and 2-Me (50μM)) and plated at 2×10^5^ cells/well in a 96 round bottomed well plate. Splenocytes were stimulated with anti-CD3 (145-2C11, 0.1μg/ml) and anti-CD28 (37.51, 1μg/ml). Cells were stained with appropriate antibodies and analysed by flow cytometry.

### Flow cytometry

For cell surface staining antibodies were incubated with cells in the dark for 30 minutes at 4°C in PBS/1% BSA. Intracellular staining was performed using Foxp3 staining buffer kit (ThermoFisher-eBioscience) according to the manufacturer’s protocol. Cells from ovarian cancer patients were incubated for 10 minutes with 10mg/ml KIOVIG (Baxalta) before staining with relevant antibodies. Cells were stained with either fixable eFluor780 Live/Dead stain (ThermoFisher-eBioscience) or aqua live/dead viability stain (ThermoFisher-Invitrogen) at 4°C in PBS.

Fluorophore conjugated antibodies against the following cell markers were used: Mouse samples – mOX40 (OX-86), CD8a (53-6.7), CD4 (GK1.5), CD3 (145-2C11), NK1.1 (PK136), FoxP3 (FJK-16s), B220 (RA3-6B2), CD11c (N418), F4/80 (BM8), CD11b (M1/70.15), CD62L (MEL-14), CD44 (IM7), KLRG1 (2F1), CD127 (A7R34), CXCR3 (CXCR3-173), CD45.2 (104), T-bet (eBio4B10) and EOMES (DAN11MAG) all from ThermoFisher-eBioscience. Anti-CD43 (activation glycoform, 1B11), Ly6C (HK1.4) and Ly6G (1A8) was purchased from Biolegend and anti-granzyme B (GB11) from ThermoFisher-Invitrogen. Isotype controls purchased from corresponding companies. H-2K^b^/SIINFEKL tetramer was prepared in house.

Healthy human samples – CD4 (RPA-T4), CD8 (SK1), hOX40 (ACT35), CD127 (eBioRDR5), Foxp3 (236A/E7), CD56 (CMSSB) and CD14 (61D3) all from ThermoFisher-eBioscience. Anti-CD3 (SK7), CD25 (M-A251) and CD19 (HIB19) were from BioLegend.

Human ovarian cancer samples and Human cells in NSG experiments - CD4 (RPA-T4), CD25 (M-A251), CD127 (HIL-7R-M21), CD8 (RPA-T8), OX40 (ACT35), and mouse IgG2a isotype,κ control (G155-178) all from BD Biosciences.

All flow cytometry experiments were performed using either a FACSCalibur, FACSCanto, FACSAria or FACSVerse machine (BD Bioscience). Analysis of data collected on the FACSCalibur was performed using Cellquest Pro (BD Bioscience) and other data was analysed using FACSDiva (V6.1.2) software (BD Bioscience) or FlowJo (BD Bioscience). Histogram overlays were produced using FCS Express (V.3) software (De Novo Software).

### OT-I Adoptive Transfer

1×10^5^ hOX40 KI^+/−^ OT-I cells were injected i.v. into hOX40 KI^+/+^ or WT C57BL/6 mice. 24 hours later 5mg ovalbumin (Sigma) and 100μg anti-hOX40 or isotype control were given via i.p. injection. Deletion was determined by harvesting spleens day 4 post i.p. injection. OT-I kinetics were monitored in the blood through SIINFEKL tetramer staining and mice were rechallenged between 6-10 weeks later with 30nM SIINFEKL i.v.

Assessment of Treg cell depletion in reconstituted NOG/SCID mice Approximately two weeks after reconstitution of NOG or NSG mice with hPBMCs, spleens were harvested and processed into a single cell suspension, followed by analysis of hOX40 expression on human T cells by flow cytometry. To examine Treg depletion, SCID or NSG mice were injected with 10×10^6^ splenocytes from reconstituted NOG or NSG mice into the peritoneal cavity 1 hour prior to injection with 10 mg/kg of depleting mAb or isotype control mAb. The peritoneal fluid was collected after 24 hours and human T cell subsets were identified by flow cytometry (see above).

### Tumour Models

E.G7 Ova and MCA-205 tumour models: 5×10^5^ tumour cells were injected subcutaneously (s.c.) into the flank of mice. Once tumours reached a certain size (E.G7:5×5-10×10mm, MCA-205: 5×5mm) mice were treated with 3×100μg shots of anti-hOX40 mAb every other day via intraperitoneal injection. For phenotyping experiments, organs were harvested, processed and analysed via flow cytometry on day 4 post final injection. Tumours were diced into smaller lumps and digested using 0.5 Units of liberase TL (Roche). For survival experiments tumour size was monitored 3 times a week and mice culled once they reached a terminal stage of the disease indicated by lump size (E.G7: 20×20mm, MCA-205: 15×15mm). Mice presenting with no tumour after treatment were rechallenged with 5×10^5^ tumour cells s.c. into the flank and again tumour size monitored every other day. Surviving mice were culled around day 100-120.

### Statistics

All results show mean ± SEM. GraphPad Prism (v7) software was used to perform statistical analysis on data and produce graphs. One way Annova with multiple comparisons (Dunnett’s, Tukey’s or Sidak’s as stated in legend) or Mann Whitney tests were used to calculate significance between groups as stated in legends. Significance on survival curves was evaluated using a Log-rank (Mantel-Cox) test. Significance shown relative to isotype control unless bar is shown. Where indicated ns = not significant, * P≤ 0.05, ** P≤ 0.01, *** P ≤ 0.001, **** P≤ 0.0001. All statistically significant pairwise comparisons are shown.

## Acknowledgements

We are grateful to the staff of the University of Southampton Biomedical Research facility for their technical support. We also thank Leon Douglas and Patrick Duriez from the ECMC/CRUK funded Protein Core Facility for making SIINFEKL tetramers

## Author contributions

JG, KH, HLS, KC, RRF, HTC, TS, MS, LM and JW performed experiments. JG, KH, HLS, TS, RRF, MS, LM and JW performed statistical analyses. JG, JW, IT, BF and MSC designed experiments. JG, JW and MSC wrote the manuscript. All authors contributed to manuscript revision and read and approved the submitted version.

## Funding

This work was supported by CRUK programme grants awarded to MJG and MSC (Award number: A20537, A24721), CRUK centre grant (Award number: A25139) and a CRUK studentship to JG and MSC from the Southampton CRUK centre (Award number: A29286).

## Conflict of interest

MSC is a retained consultant for BioInvent International and has performed educational and advisory roles for Baxalta and Boehringer Ingleheim. He has received research funding from Roche, Gilead, Bioinvent International and GSK. BF, IT, LM and MS are employees of Bioinvent International.

