## Supplementary Figures 1-5 for "Domain Binding and Isotype Dictate the Activity of Anti-human OX40 Antibodies"

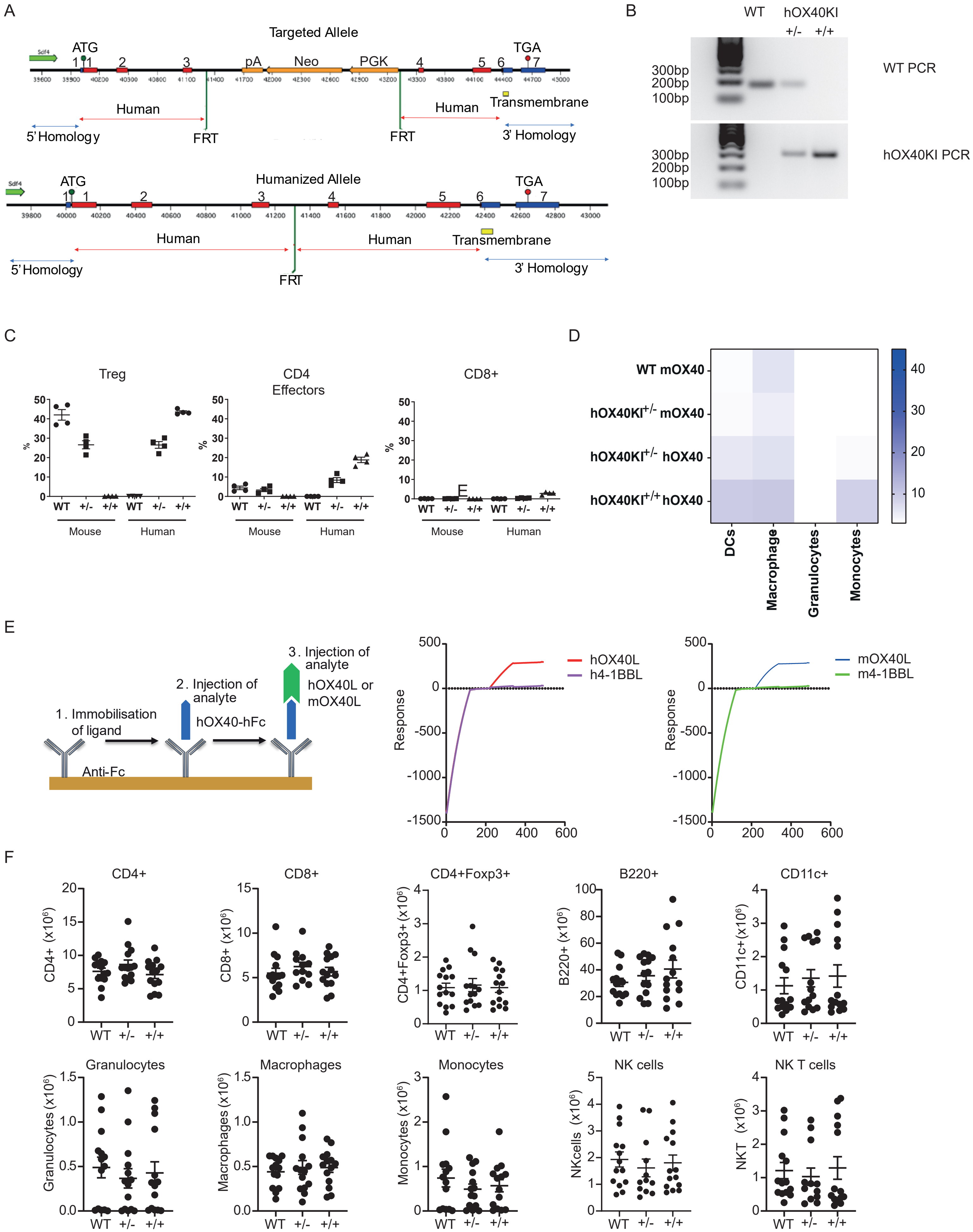


Supplementary Figure 1. hOX40 expression in hOX40KI mice is dose dependent. A. Schematic of hOX40 chimeric receptor construct used to generate the hOX40KI mice. B. PCR showing the genotyping of WT, hOX40KI$^{+/-}$ and hOX40KI$^{+/+}$ mice. C. Expression of mouse and human OX40 on Treg (left panel), CD4 effectors (middle panel) and CD8 T cells (right panel) isolated from WT, hOX40KI $^{+/-}$ and hOX40KI$^{+/+}$ n=4. D. Heat map showing expression levels on myeloid populations. E. SPR analysis of OX40 L binding to hOX40. Left panel shows schematic, h4-1BBL and m4-1BBL were used as negative controls. Middle panel shows hOX40L (red) and h4-1BBL (purple) binding and right panel shows mOX40L (blue) and m4-1BBL (green) F. Lymphocyte and Myeloid populations in WT, hOX40KI$^{+/-}$ and hOX40KI$^{+/+}$ mice (6-10 weeks age, n=14).

Supplementary Figure 2

A

| Antibody | Parental Isotype | Domain binding | ka (M-1·s-1) | kd (s-1) | KD (M) |
|---|---|---|---|---|---|
| SAP 28-2 | mIgG1 | 1 | $4.2 \times 10^{4}$ | $2.57 \times 10^{-4}$ | $6.1 \times 10^{-9}$ |
| SAP 28-3 | mIgG1 | 2 | $2.5 \times 10^{5}$ | $0.46 \times 10^{-4}$ | $1.84 \times 10^{-10}$ |
| SAP 15-3 | mIgG1 | 2 | $14.3 \times 10^{4}$ | $4.68 \times 10^{-4}$ | $3.3 \times 10^{-9}$ |
| SAP 9 | mIgG2a | 3 | $50.9 \times 10^{4}$ | $3.98 \times 10^{-4}$ | $7.82 \times 10^{-10}$ |
| SAP 25-29 | mIgG1 | 4 | $2.56 \times 10^{4}$ | $1.31 \times 10^{-4}$ | $5.1 \times 10^{-9}$ |
| SAP 29-50 | mIgG1 | 4 | $13.8 \times 10^{4}$ | $0.62 \times 10^{-4}$ | $4.49 \times 10^{-10}$ |
| SAP 29-23 | mIgG2a | 4 | $29 \times 10^{4}$ | $1.81 \times 10^{-4}$ | $6.24 \times 10^{-10}$ |

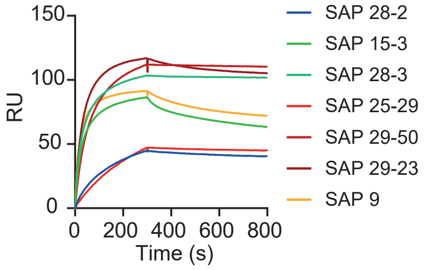


B

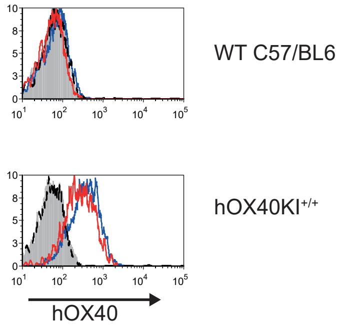


C

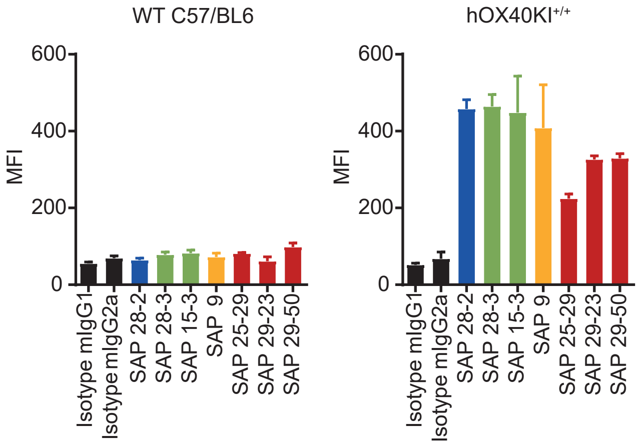


D

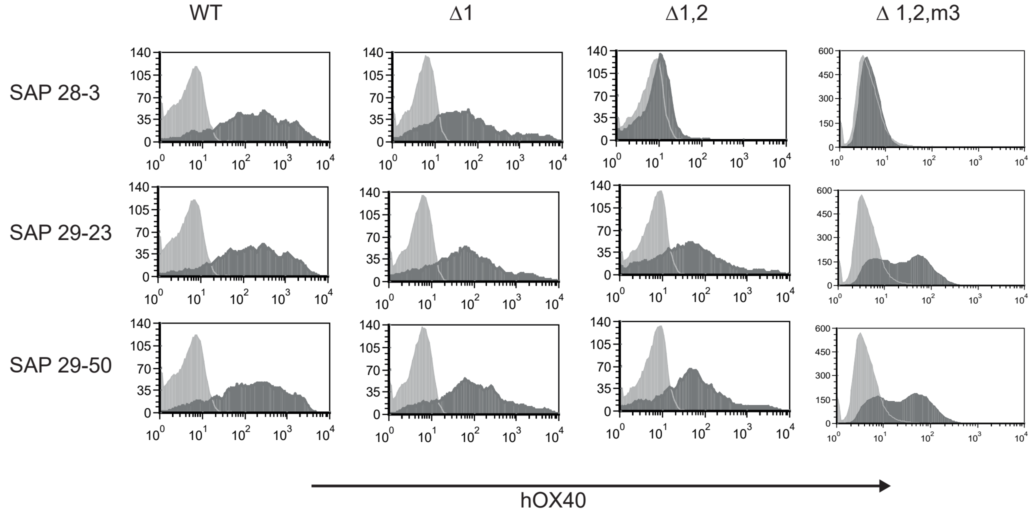


E

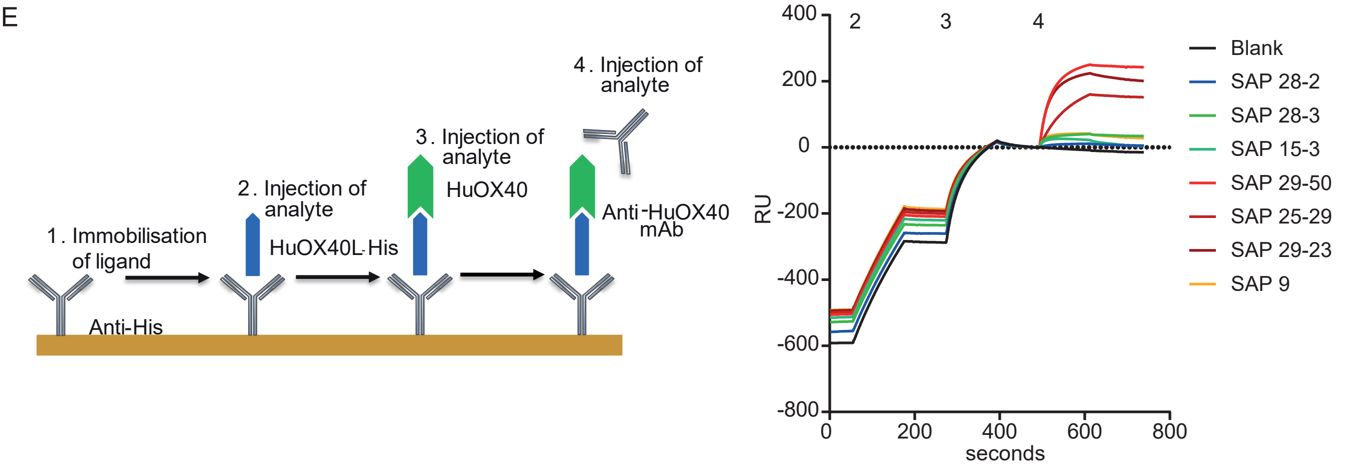


Supplementary Figure 2. Characterisation of anti-hOX40 mAb. A. SPR analysis of anti-hOX40 mAb binding and affinity. B. Binding of anti-hOX40 mAb SAP28-2 mIgG1(blue line) and SAP 29-23 mIgG2a (red line) to activated Tregs (cells activated with αCD3 (0.1μg/ml) and αCD28 (5μg/ml) for 24 hours) from WT (top panel) or hOX40KI$^{+/+}$ mice (bottom panel). Isotype controls shown as filled grey histogram (mIgG1) and black dotted line (mIgG2a). C. Binding of all parental anti-hOX40mAb to activated Tregs (cells activated as in B) isolated from WT (left bar chart) or hOX40KI$^{+/+}$ mice (right bar chart) n=3. D. Binding of FITC-labelled anti-hOX40 mAb to hOX40 domain mutants. E. Schematic and full SPR analysis of anti-hOX40mAb binding in the presence of hOX40L.

Supplementary Figure 3

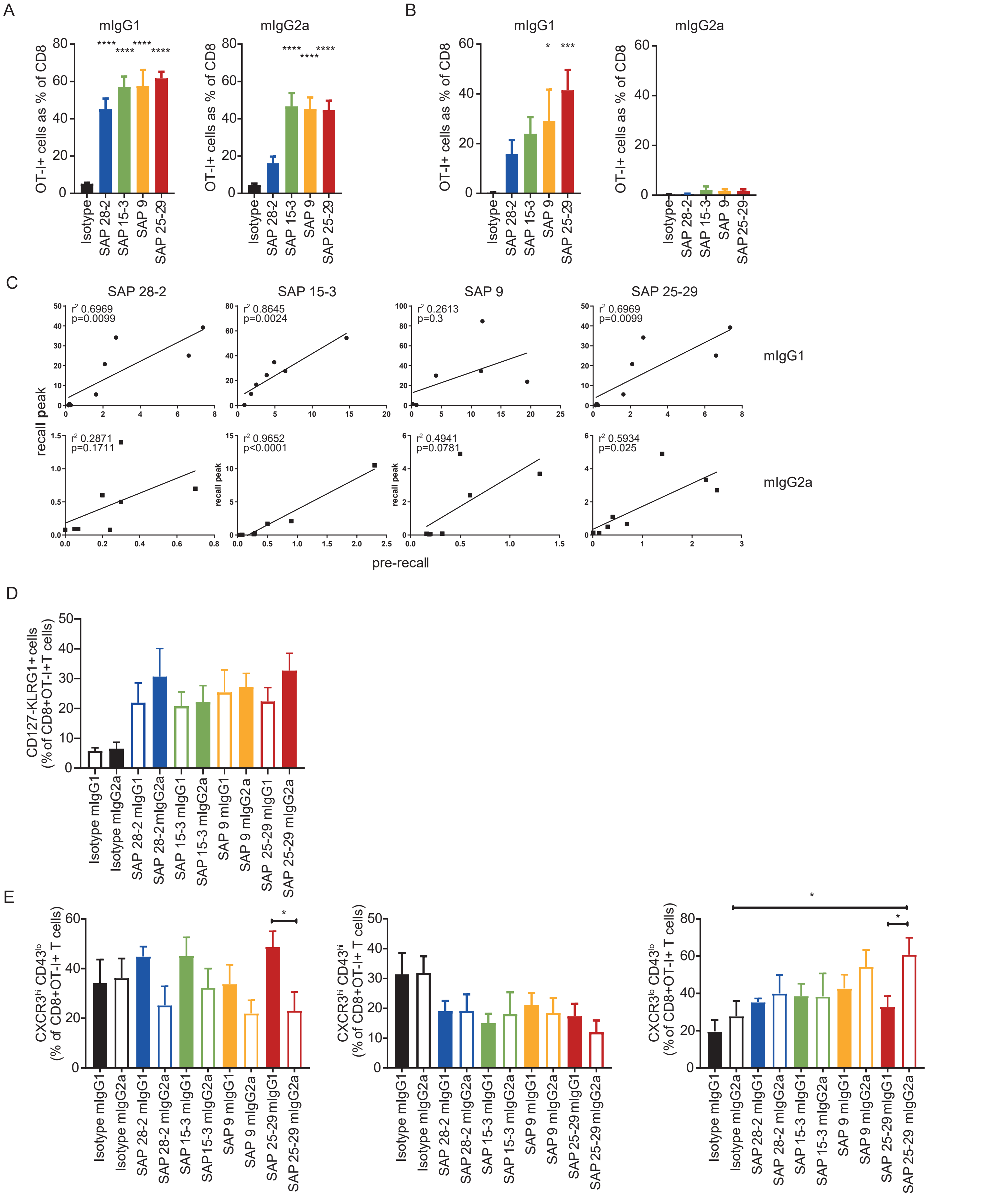


Supplemental Figure 3. anti-hOX40 mIgG1 act agonistically *in vivo*. A. Primary peak response of OT-I expansion in blood to anti-hOX40 mIgG1 (left panel) or mIgG2a (right panel) (n=6-8, pooled from 2 independent experiments). B. Recall peak following SIINFEKL rechallenge of mice previously treated with anti-hOX40 mIgG1 (left panel) or mIgG2a (right panel) (n=6-8, pooled from two independent experiments). C. Correlation graphs between recall response and pre-recall OT-I levels in anti-hOX40 mIgG1 (top row) and mIgG2a (bottom row) treated mice. n=6-8, pooled from two independent experiments. D. Analysis of OT-I SLECs CD127-KLRG1+ in the blood at D18 (n=8). E. CXCR3 and CD43 analysis of OT-I in the blood pre-rechallenge with SIINFEKL peptide (n=7-8 pooled from 2 independent experiments). CXCR3hiCD43lo (left panel), CXCR3hiCD43hi (middle panel) and CXCR3loCD43lo (right panel). *p<0.05 Sidak's test.

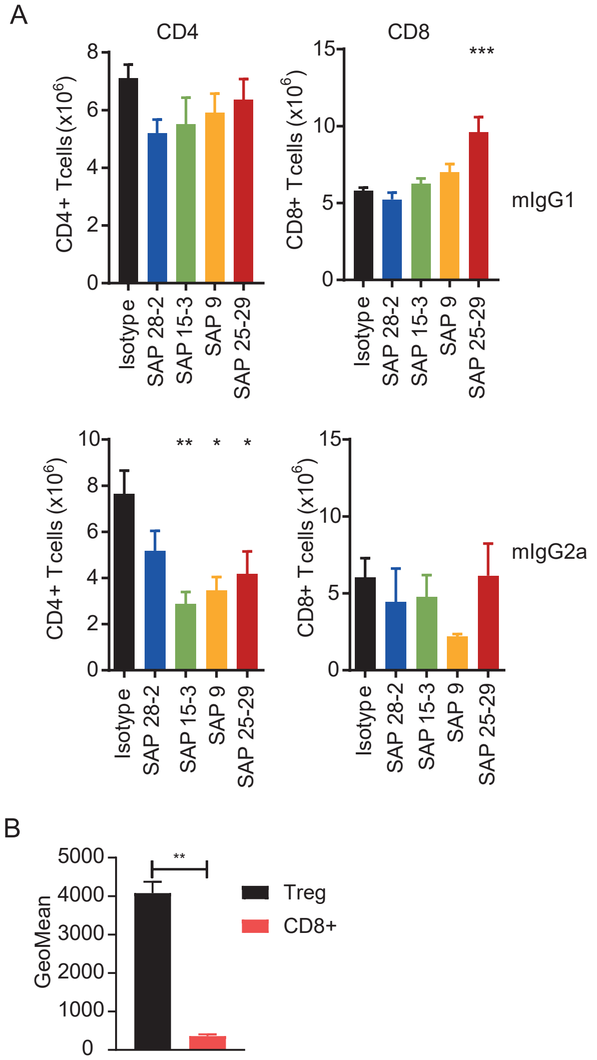


Supplemental Figure 4. Isotype of anti-hOX40 mAb influences effector function. A. CD4 and CD8 T cell numbers assessed on Day 4 in splenocytes isolated from hOX40KI$^{+/+}$ mice receiving 1x10$^5$ hOX40KI$^{+/-}$OT-I followed by 100μg anti-hOX40 mAb as either a mIgG1 or mIgG2a (n=7-8, pooled from 2 independent experiments). ** p<0.01, * p<0.05 Dunnett's test. B. hOX40 expression on Treg and CD8+ T cells isolated from NOG mice 2 weeks post injection of hPBMCs (n=5, representative of 2 independent experiments). ** p<0.01 Mann Whitney U test.

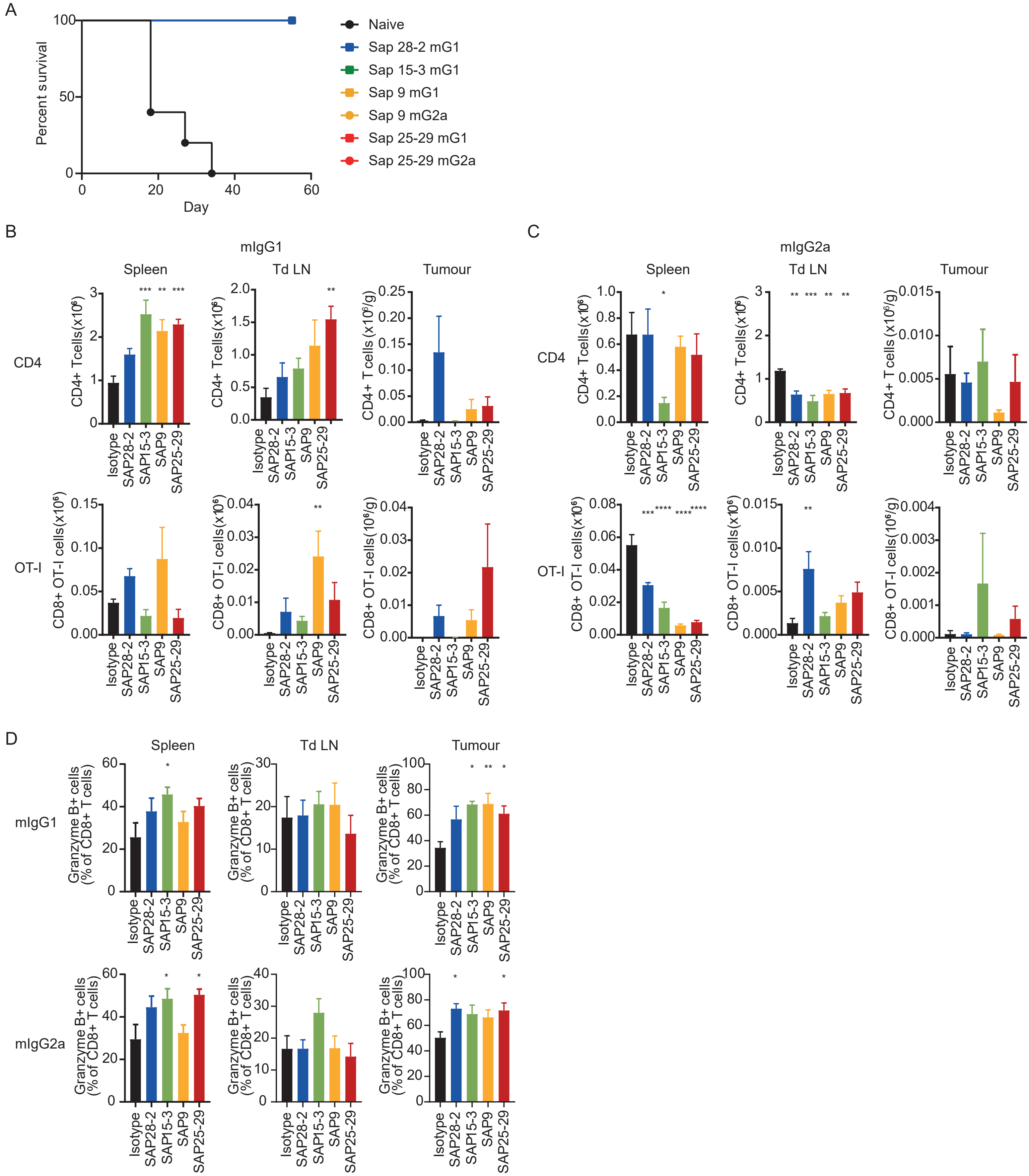


Supplementary Figure 5. hOX40mAb are therapeutic as both mIgG1 and mIgG2a. A. Survival graphs of mice rechallenged with E.G7 lymphoma cells shown alongside naive recipients. n=1-5 representative of 2 independent experiments. B and C. Assessment of CD4+ (top rows) and CD8+OT-I (bottom rows) T cell populations in Spleen (left panels), Tumour draining lymph node (middle panels), and Tumour (right panels) isolated 24 hours post second mAb dose either as a mIgG1 (B) or mIgG2a (C). n=3-5, representative of 2 independent experiments. D. Analysis of Granzyme B expression on CD8+T cells isolated from Spleen (left panels), Tumour draining lymph node (middle panels) and Tumour (right panels) from mice treated with either mIgG1 (top row) or mIgG2a (bottom row). Data pooled from 2 independent experiments (n=8-9). ****p<0.0001, *** p< 0.001 ** p<0.01 * p<0.05 Log-rank (Mantel-Cox) for survival graphs A and B and Dunnett's multiple comparison test for B-D.
